# Delayed puberty, gonadotropin abnormalities and subfertility in male *Padi2/Padi4* double knockout mice

**DOI:** 10.1101/2022.02.09.479603

**Authors:** Kelly L. Sams, Chinatsu Mukai, Brooke A. Marks, Chitvan Mittal, Elena Alina Demeter, Sophie Nelissen, Jennifer K. Grenier, Ann E. Tate, Faraz Ahmed, Scott A. Coonrod

## Abstract

**Background:** Peptidylarginine deiminase enzymes (PADs) convert arginine residues to citrulline in a process called citrullination or deimination. Recently, two PADs, PAD2 and PAD4, have been linked to hormone signaling *in vitro* and the goal of this study was to test for links between PAD2/PAD4 and hormone signaling *in vivo.*

**Methods:** Preliminary analysis of *Padi2* and *Padi4* single knockout (SKO) mice did not find any overt reproductive defects and we predicted that this was likely due to genetic compensation. To test this hypothesis, we created a *Padi2/Padi4* double knockout (DKO) mouse model and tested these mice along with wild-type FVB/NJ (WT) and both strains of SKO mice for a range of reproductive defects.

**Results:** Controlled breeding trials found that male DKO mice appeared to take longer to have their first litter than WT controls. This tendency was maintained when these mice were mated to either DKO and WT females. Additionally, unsexed 2-day old DKO pups and male DKO weanlings both weighed significantly less than their WT counterparts, took significantly longer than WT males to reach puberty, and had consistently lower serum testosterone levels. Furthermore, 90-day old adult DKO males had smaller testes than WT males with increased rates of germ cell apoptosis.

**Conclusions:** The *Padi2/Padi4* DKO mouse model provides a new tool for investigating PAD function and outcomes from our studies provide the first *in vivo* evidence linking PADs with hormone signaling.

## BACKGROUND

Peptidylarginine deiminases (PADs or PADIs) are a family of calcium-dependent enzymes that post-translationally convert positively-charged arginine residues to neutrally-charged citrulline in a process called citrullination or deimination. The loss of charge on target arginine residues following citrullination can significantly alter the target protein’s tertiary structure and affect protein-protein and protein-DNA/RNA interactions (1). There are five PAD family members (PAD1-4 and 6), with each of the isoforms showing some degree of substrate and tissue specificity (2). Functional roles for PADs in mammalian physiology and pathology are diverse and include cellular differentiation, nerve growth, apoptosis, inflammation, early embryonic development, and gene regulation (3).

While PAD activity is most closely associated with autoimmune disease (4), previous studies have found that PAD2 and PAD4 are expressed in reproductive epithelial tissues (ie. uterus, luminal breast tissues, endometrium, leydig and sertoli cells) and that their expression is regulated by steroid hormone signaling (5–8). Steroid hormones are essential mediators of many physiological events including reproduction and development (9). These hormones carry out their effects by binding to their cognate nuclear receptors which, in turn, bind to hormone response DNA elements and regulate transcription via interaction with a wide range of co-regulators and the basal transcriptional machinery. Given the potential links between PADs and steroid hormones and that histones were found to be targeted by PAD4 for citrullination (10), we previously tested whether PAD2 and PAD4 may play a role in estrogen signaling in breast cancer cells by facilitating ER target gene expression via histone citrullination at ER binding sites. Results from our studies found that PAD4 is recruited to the TFF1 promoter (a canonical ERα binding site) in MCF7 cells following E2 treatment leading to citrullination of histone H4 arginine 3 (11). Additionally, we found that PAD2 interacts with ER in an estrogen-dependent manner and that estrogen (E2) treatment activates PAD2 leading to citrullination of histone H3 arginine 26 (H3Cit26) at ER binding sites (12,13). We then demonstrated by ChIP-Seq that ~95% of ER binding sites throughout the genome are citrullinated at H3Cit26 within 5-10 minutes of E2 treatment (14). We also found that PAD inhibitors potently blocked ER binding at Estrogen Response Elements (EREs) and prevented ER target gene activation (13).

Regarding a potential role for PADs in hormone signaling in males, another group recently found that PAD2 plays a role in androgen receptor (AR) signaling by facilitating nuclear import of AR and promoting AR target gene activation via histone citrullination at AR binding sites in castration resistant prostate cancer cell lines (CRPC; LnCaP and VCaP) (12). This finding suggests that the role of PADs in regulating hormone-mediated transcription may be broader than we had previously realized.

Recent studies have already begun to document the presence of PAD2 in the testis. For example, Tsuji-Hosokawa et al. showed that PAD2 appears to play an important role in testis development and is specifically expressed in fetal Sertoli cells, and that this expression is regulated by SOX9 a critical mediator of sex determination (15). Another recent study used PCR, western blotting, and IHC to show that *Padi2* is expressed in sperm during the first wave of spermatogenesis and appears to localize to the acrosomal region of the sperm head (16). There are currently no reports of PAD4 localization to the testes, however, IHC analysis of the testis from the Human Protein Atlas (proteinatlas.org) does show anti-PAD4 staining (moderate intensity) specifically in Leydig cells (6). Importantly, to date, there have not been any published reports linking PADs to hormone signaling in animal models and this lack of *in vivo* data has likely limited research into this area.

To address this gap in knowledge, we began testing for links between PADs and hormone signaling *in vivo* by determining whether *Padi2* and *Padi4* single knockout (SKO) mouse lines displayed overt reproductive defects. Initial observations with these mice found that both *Padi2* and *Padi4* SKO mice appear normal with no overt reproductive defects. Given that the amino acid sequence in PAD family members is 59-70% identical, and that *Padi* genes are closely clustered together at chromosome 1p35-36 in humans and chromosome 4E1 in mice (3,17), we predicted that the lack of a conspicuous phenotype in the SKO lines was due to genetic compensation by other *Padis.* In order to test this hypothesis, we generated a *Padi2/Padi4* double knockout mouse (DKO) line by deleting *Padi4* from our existing *Padi2* knockout (P2KO) mouse line using CRISPR/CAS9 technology. We demonstrate that, as opposed to the SKO lines, the *Padi2/Padi4* DKO mice display a range of reproductive defects. Outcomes from our studies focusing on the DKO males show that they display a reduced pup and weanling weight, delayed puberty, reduced testis size, increased rates of apoptosis during spermatogenesis, and altered circulating hormone levels. Our results provide the first *in vivo* evidence that PADs play a role in male reproduction, potentially via their role in hormone signaling.

## MATERIALS AND METHODS

### Animals

All mice used in this work were on an FVB/NJ background and were maintained as inbred homozygous lines. The mice were housed in an AAALAC certified facility, with 12 hours on/off light cycle, temperature control and food and water ad libitum. All procedures were approved by Cornell University’s Animal Care and Use Committee (IACUC) and in compliance with the US National Research Council’s <u>Guide for the Care and Use of Laboratory Animals</u> (18). Breedings were recorded and lineages tracked with Softmouse Colony Management Software (19). Euthanasia was performed by CO_2_ inhalation in accordance with American Veterinary Medical Association approved practice.

### Generation of transgenic single knockout mice

*Padi2-null* mice (FVB/NJ background) were generated using gene trap technology at the Texas A&M Institute for Genomic Medicine (20). For the gene trap, 163 nucleotides of exon 1 in *Padi2* (from the ATG initial codon to intron 1-2) were replaced by the LacZ and Neomycin coding sequence. These mice are genotyped in a multiplex mixture using *Padi2* WT and KO specific primers (Table 1).

**Table 1.**
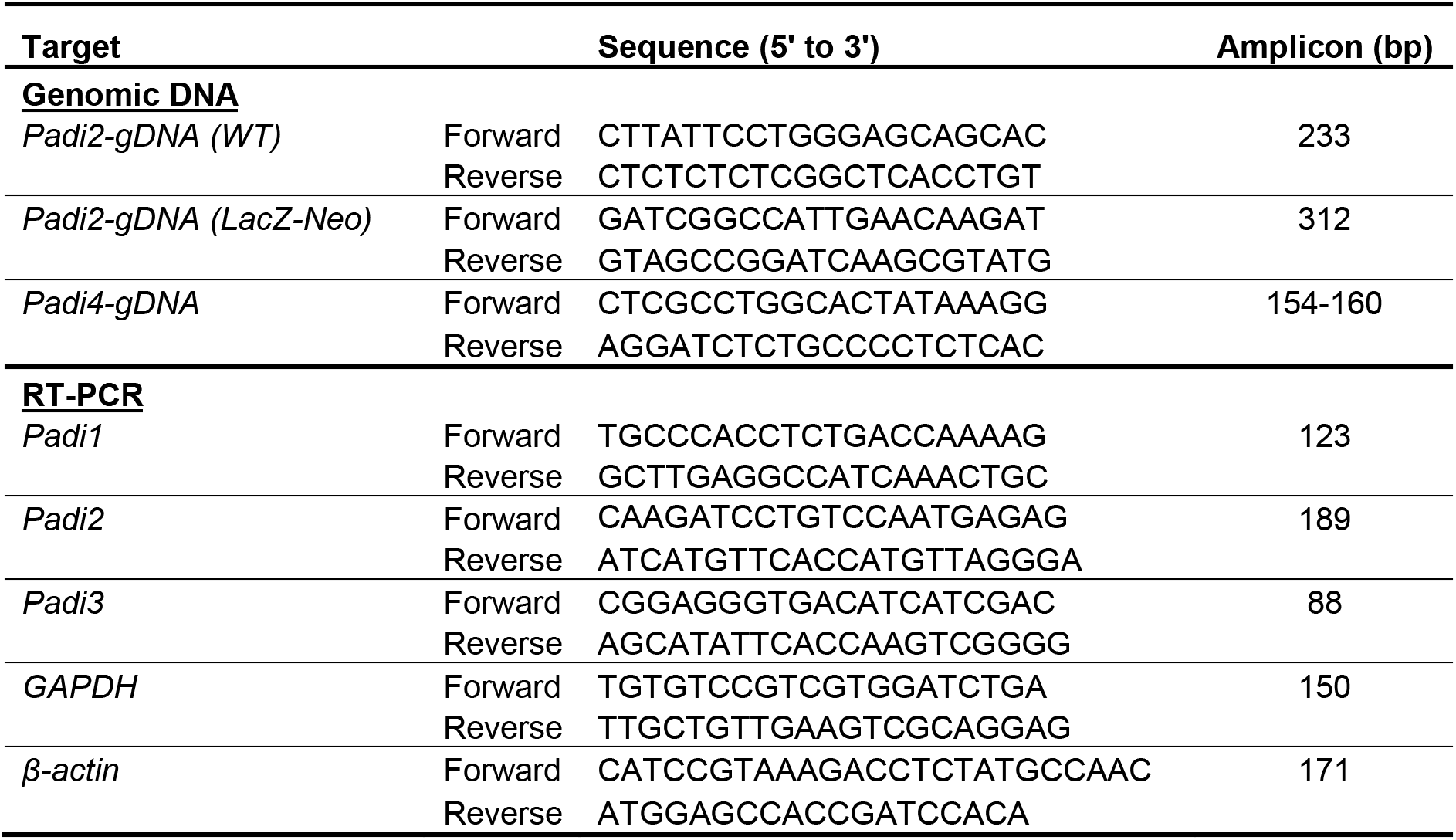
Primers used for genotyping and authentication of DKO mice.

*Padi4-null* mice (FVB/NJ background) were generated using Regeneron’s VelociGene technology. The genomic sequence of *Padi4* from the ATG initial codon to approximately 100bp downstream of the TGA stop codon was replaced in frame (with respect to the *Padi4* initiation codon) with a Lox-flanked LacZ and neomycin coding sequence. These mice are genotyped using *Padi4* primers shown in Table 1.

### Generation of Padi2/Padi4 double knockout mice

*Padi2-null* mice were used for IVF and CRISPR/Cas9 RNA injection was performed at Cornell’s Stem Cell & Transgenic Core Facility. Cas9 mRNA and *Padi4* sgRNA targeting Exon 1 of the *Padi4* gene was injected into the pronucleus of *Padi2* KO embryos and the embryos were cultured to the two-cell stage and transferred into pseudopregnant recipient FVB/NJ female mice (21,22). Thirty-nine founders were produced and 13 edited founders were then identified by heteroduplex PCR (23). Briefly, genomic DNA was extracted from punched ear tissue and evaluated by PCR using the *Padi4* primer set noted above. Standard PCR conditions for GoTaq (Promega) were used as follows: 95°C for 5 min; 94°C for 30 s, 59°C for 30 s, 72°C for 30s for 35 cycles; 72°C for 5 min followed by 5 minutes of denaturation at 95°C. PCR products were loaded onto a 12% polyacrylamide gel. Positive heteroduplex bands were purified and subjected to TA cloning for Sanger sequencing. All 13 edited founders were found to be heterozygous mutants, with 7 founders being identified as having a frameshift mutation either by insertion or deletion. One heterozygous founder with a four-nucleotide deletion was back-crossed to the background *Padi2* knockout mice, and their heterozygous offspring were then paired to create a homozygous DKO line (Figure 1).

**Figure 1.**
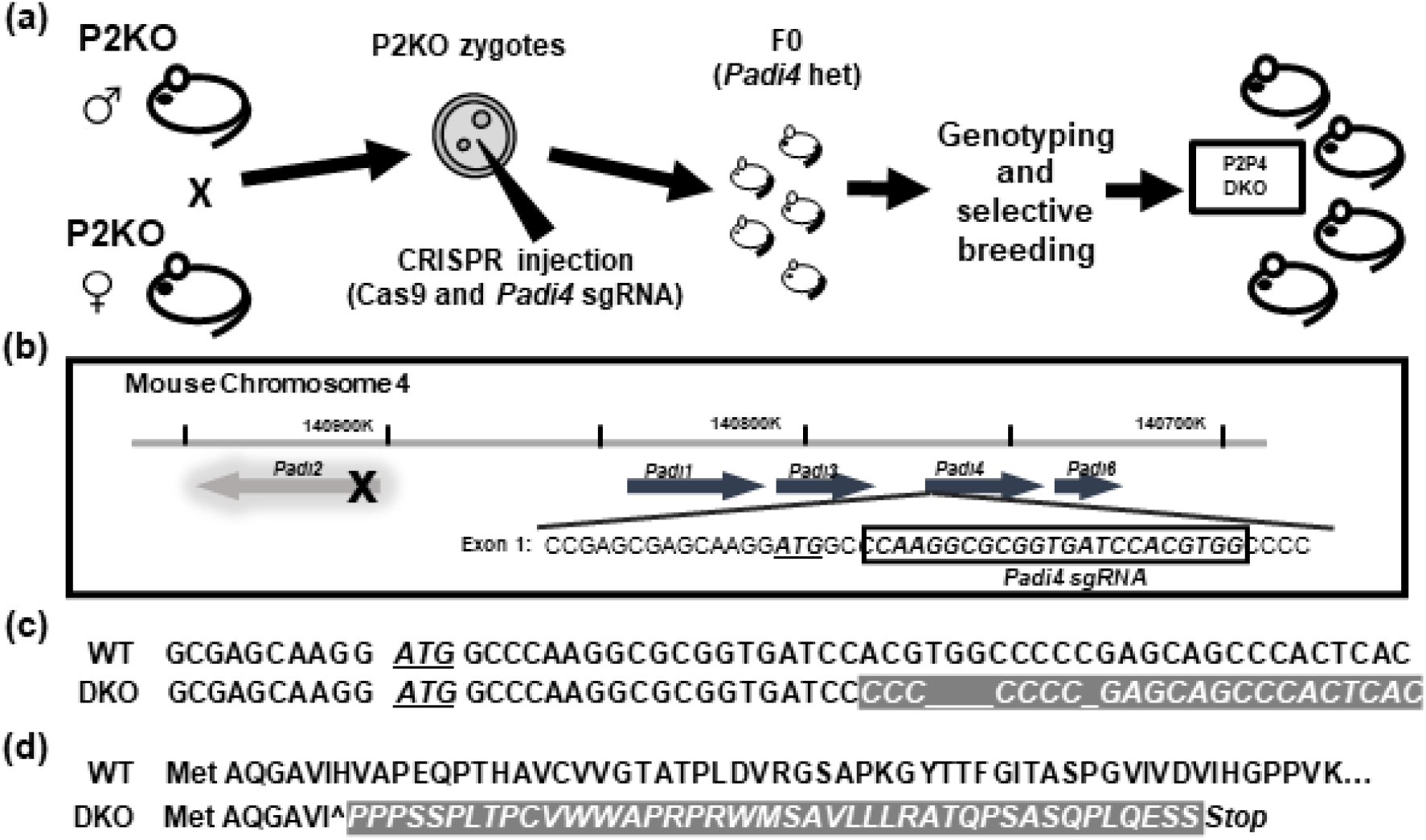
Design of *Padi2/Padi4* double knockout mice. (a) Schematic of the CR1SPR protocol using *Padi2* KO zygotes. (b) Design and sequence of the guide RNA targeting Exon 1 of the *Padi4* gene. (c) Sanger sequencing confirmed afour-nucleotide deletion in the DKO *Padi4* Exon 1, resulting in (d) a frame-shift mutation and premature stop codon in the predicted amino acid sequence.

### Authentication of Padi2/Padi4 double knockout mice

#### RT-PCR of Padi transcripts

Total RNA was extracted from mouse salivary glands using Tri-Reagent (Molecular Research Center #TR118) and cDNA was created with a High Capacity RNA-to-cDNA Kit (Applied Biosystems #4387406) according to manufacturers’ instructions. PCR amplification was accomplished under standard cycling conditions with GoTaq (Promega #M3001) using primers listed in Table 1. PCR products were run on a 2% agarose gel to confirm the presence or absence of *Padi* transcripts (Figure 2a,d; Additional File 1).

**Figure 2.**
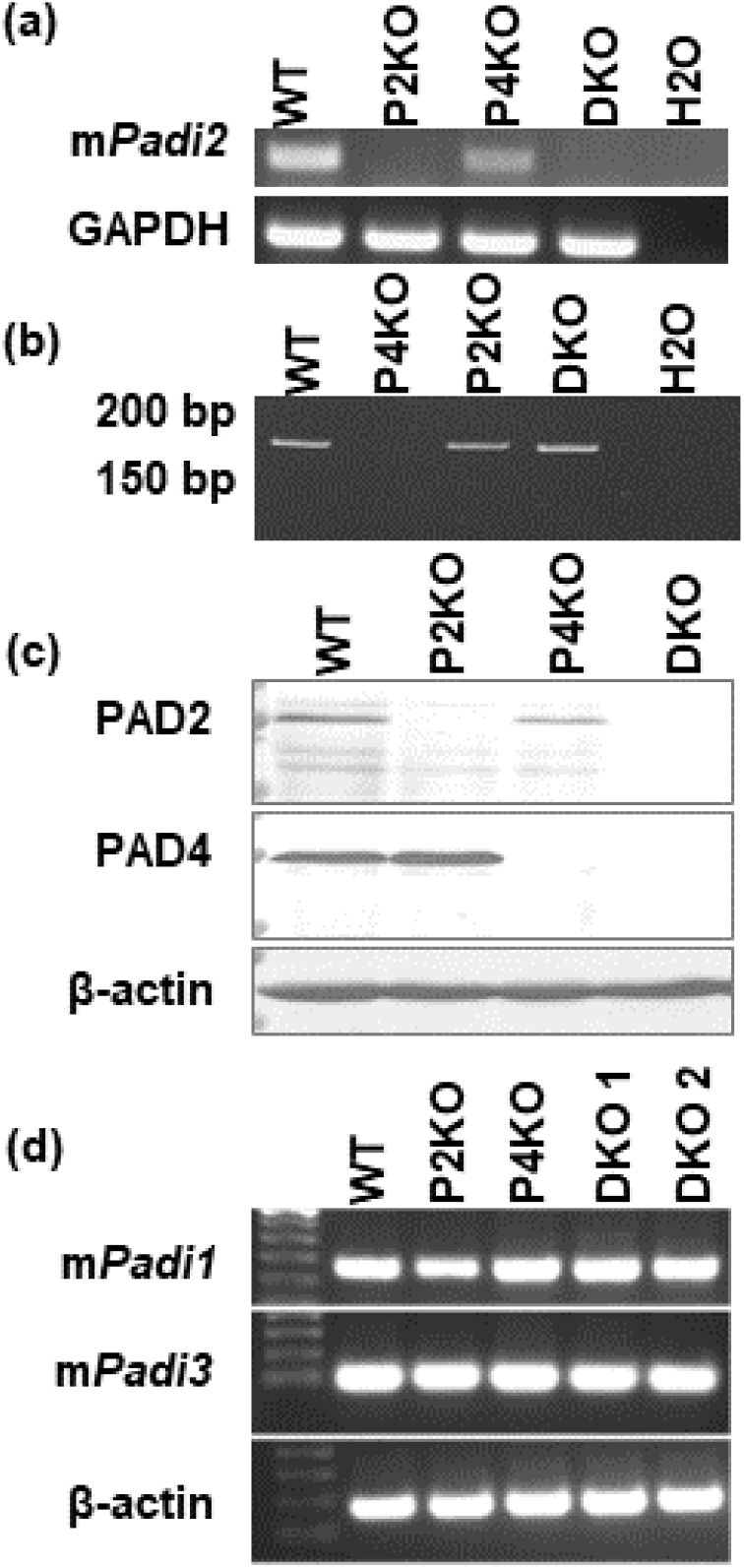
Authentication of the *Padi2/Padi4* double knockout. (a) RT-PCRof salivary gland tissue shows that the DKO mice lack Padi2 transcripts. (b) PCR analysis of genomic DNA (15% polyacrylamide gel) confirms a 4bp deletion within the *Padi4* gene of DKO mice. (c) Western blot analysis of spleen lysate confirms that DKO mice lack both PAD2 and PAD4 mature proteins. (d) RT-PCR of testicular tissue confirms transcription of both *Padi1* and *Padi3* is unaffected by the deletion of nearby genes.

#### PCR of Padi4 locus

Genomic DNA was isolated from ear-punch tissue samples and the *Padi4* locus was amplified under standard GoTaq conditions using primers listed in Table 1. PCR products were run on a 2% Agarose gel to confirm the presence or absence of the *Padi4* gene (Figure 2b).

#### *Western blot for* PAD2 *and* PAD4 *protein products*

Tissue lysate was prepared from splenic tissue and resolved by SDS-PAGE followed by transfer to a PVDF membrane. The membranes were incubated overnight with primary antibodies diluted in 1% BSA-TBST at 4°C using the following concentrations: anti-PAD4 (1:1000; Abcam; ab214810), anti-PAD2 (1:1000, Proteintech, 12110-1-AP) and anti-actin (1:1000, Abcam, ab-8227). The primary antibodies were detected with HRP-conjugated secondary antibodies and were exposed to ECL reagents to determine the presence or absence of PAD2 and PAD4 proteins (Figure 2c).

### Breeding Trials

Seven-week old virgin mice were paired in the following ‘male x female’ matings: DKOxDKO, DKOxWT, WTxDKO or WTxWT (n=5-6 pairs per group). Pairs were monitored daily for 63 days and birth dates, litter sizes, sex ratios, 2-day old pup weights, and weanling weights were recorded. Statistical analyses were carried out using Microsoft Excel and Social Science Statistics calculators for unpaired Student’s t-test, ANOVA, *χ*2 and Goodness of Fit tests with significance set at *P* = 0.05 (24).

### Growth and development

Male weanlings were housed singly (to avoid intra-cage dominance issues), weighed and monitored daily for preputial separation as previously described (25). Growth curves are provided in Supplemental Figure S2 (Additional File 2). Mice were euthanized for tissue harvest at ages 46, 48, 50 and 90 days old. At each of these time points, five mice were euthanized by CO_2_ inhalation and blood was collected by cardiac puncture, allowed to clot at room temperature for 90 minutes, spun at 2000g for 15 minutes, and the serum was snap frozen and saved for hormone analysis. Serum testosterone, Follicle Stimulating Hormone (FSH), and Lutenizing Hormone (LH) levels were quantified in duplicate by ELISA at the University of Virginia Ligand Assay Core. Testes, epididymis, vas-deferens, seminal vessicles and spleen were harvested and one testis was immediately snap frozen for RNA analysis. The remaining tissues were fixed in 10% formalin for preliminary histological evaluation. Results were analyzed with unpaired t-tests with significance set at *P* = 0.05.

### Histological evaluation

Testis, epididymis, vas deferens, seminal vesicle and splenic tissues were collected at

90 days from 5 animals from each of the following strains: Wild-type (WT, FVB/NJ), P2KO, P4KO, and DKO. Collected tissues were fixed in 10% formalin solution, embedded in paraffin, sectioned at 5μm, and stained with hematoxylin and eosin (H&E). Initial evaluation revealed epididymis, vas deferens, seminal vesicles, and spleen within normal limits based on previously published observations (26,27), therefore, only testis sections were evaluated further. The noted changes were scored on a scale of 0 to 3 according to the following criteria: 0 = absent to affecting less than 5% of the tissue; 1 = mild change, affecting 5-25% of the tissue; 2 = moderate change, affecting 25-50% of the tissue; 3 = severe change, affecting >50% of the tissue (Table S1, Additional File 3).

### RNA sequencing

Mouse testes were dissected from four-month old mice, twoper strain (WT, P2KO, and DKO), and snap-frozen immediately following euthanasia. Total RNA extraction was performed using TRI reagent (Molecular Research Center, Inc. OH) according to the manufacturer’s protocol. The RNA concentration and purity were measured by Nanodrop and Qubit (Thermo Scientific, USA).

RNA-seq experiments were performed in the Transcriptional Regulation & Expression Facility (TREx) at Cornell University. RNA integrity was confirmed on a Fragment Analyzer (Agilent) and RNAseq libraries were prepared using the NEB Ultra II Directional RNA library Prep kit (New England Biolabs), with 1 ug total RNA input and initial poly(A) selection. Illumina sequencing was performed on a NovaSeq to generate at least 20M 2×150 paired end reads per sample. The raw fastq reads were first processed with the Trim Galore package to trim for low quality reads, noisy short fragments, and adapter sequence (28). The filtered reads were then aligned to the GRCm38 reference genome (mm10) with ENSEMBL annotations via STAR using parameters [--outSAMstrandField intronMotif, --outFilterIntronMotifs RemoveNoncanonical, --outSAMtype BAM SortedByCoordinate, --quantMode GeneCounts] (29). Differential gene expression analysis was performed by DESeq2 (30,31), with an FDR cut-off of 0.05 and a Benjamini-Hochberg adjusted *P* value (*p-adj*) < 0.05 to establish significance. A heatmap and hierarchical clustering dendrogram of differentially expressed genes was produced using Morpheus (32). Venn diagrams were produced with BioVenn (33).

Gene ontology classification analysis was performed using DAVID v6.8, a web-based bioinformatics resource for functional genomics analysis to identify biological processes, molecular functions, and cellular components (34,35). Two hundred and forty-six genes were identified as uniquely upregulated in DKO testes compared to WT. Two hundred and thirty-eight of these genes were identified in the DAVID database and used for subsequent analysis.

## RESULTS and DISCUSSION

### Generation of Padi2/Padi4 double knockout mice

Our strategy for generating the *Padi2/Padi4* DKO mouse line is shown in Figure 1a. *Padi2-null* males and females were bred to generate *Padi2-null* zygotes and these embryos were then injected with a *Padi4* sgRNA construct and transferred to wild type (FVB/NJ) recipient females to generate founder pups. The *Padi4* sgRNA construct was designed to target Exon 1 of the *Padi4* gene (Figure 1b). In total, 39 founders were born, with 13 CRISPR-edited founders being identified by heteroduplex PCR, subcloning, and Sanger sequencing. All of the edited founders were heterozygous mutants, with seven of these founders having a frame-shift mutation. Following back breeding to P2KO, crossing the heterozygous F1 offspring, and genotyping successive generations, one homozygous DKO strain was established and selected for further characterization. This mutant mouse line contained a four-nucleotide deletion in Exon 1 of *Padi4* (Figure 1c) and a predicted premature stop codon (Figure 1d).

Validation of the DKO mutant line is shown in Figure 2. We first confirmed that P2KO and DKO mice lacked *Padi2* transcripts by RT-PCR analysis of salivary gland (Figure 2a). We next tested whether the *Padi4* sgRNA targeted site in DKO mice contained a deleted sequence by performing PCR analysis of genomic DNA and then separating the DNA fragments on high percentage polyacrylamide gels. Results showed that the DKO amplicon displayed a reduced molecular weight when compared to the P2KO and WT amplicons (Figure 2b), suggesting that DKO mice contained the targeted deletion, as was shown by Sanger sequencing of the *Padi4* locus (Figure 1c). As expected, no amplicon was observed in P4KO mice. The above results suggested that DKO mice lacked both *Padi2* and *Padi4* transcripts. We then performed Western blot analysis of WT, P2KO, P4KO and DKO spleen lysates to test if the DKO mice also lacked PAD2 and PAD4 protein. We probed the protein lysates with anti-PADI2 antibodies and found that, while the antibody was reactive with appropriately sized ~75 kDa bands in the WT and P4KO lysates, the antibody did not react with similarly sized bands in the P2KO and DKO mice (Figure 2c, Additional File 1). Likewise, the anti-PADI4 antibody was reactive with ~75 kDa bands in the WT and P2KO lysates but was not reactive with similarly sized bands in the P4KO and DKO lysates (Figure 2c, Additional File 1). These results indicate that the *Padi2/Padi4* DKO strain does not express mature PAD2 and PAD4 proteins. Lastly, we found by RT-PCR that the transcripts of *Padi1* and *Padi3* continue to be expressed in our DKO mice, indicating that the observed phenotype was due to the disruption of *Padi2* and *Padi4* and that the other *Padi* loci remain intact (Figure 2d).

### Effect of Padi2/Padi4 DKO on fertility parameters

In order to begin testing for links between PADs and reproductive function, we first performed a controlled breeding trial to test whether a range of reproductive parameters were affected by DKO. Seven-week old mice were paired in 4 mating categories (WTxWT, DKOxDKO, DKOxWT and WTxDKO) and monitored daily for 63 days. Results showed that, although not statistically significant, DKOxDKO pairs tend to lag behind WTXWT pairs in the time taken to have their first litter (Figure 3a). To identify which sex may be responsible for this reproductive delay, we bred DKO males with WT females and DKO females with WT males, and the trend towards decreased fertility remains more apparent for DKO males than for females (Figure 3a).

**Figure 3.**
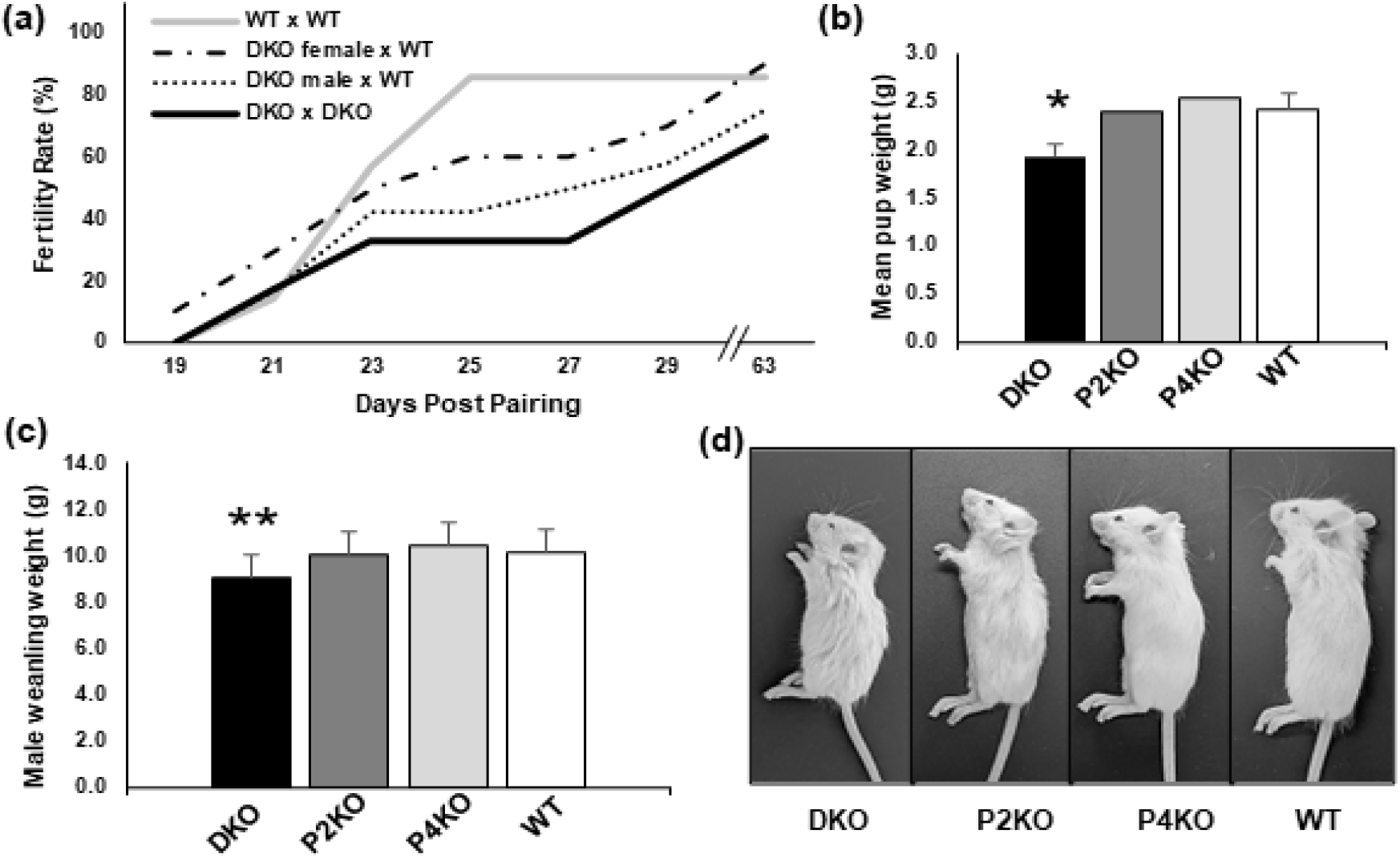
Breeding phenotype of *Padi2/Padi4* double knockout mice. (a) In controlled breed i ng tri al s. DKO paiis (n=6) took I ong er th an WT paiis (n=7) to h ave thei r first I itter. When paired with WT mates, DKO males (n=12) took longer than DKO females (n=10) to have their first litter and had fewer pup-producing pairs (data trending, but not significant, one-tailed Fisher’s Exact Test. *P*= 0.19). (b) Unsexed pups from the breedingtrial litters were weighed at two days post-pa rtum, and showed that DKO pups were smaller than pups of either SKO strain, and significantly smaller than WT pups (one-tailed t-test, \**P*= 0.004). (c) Male DKO are significantly smaller than both SKO and WT mice at weaning (ANOVA. p=0.02. post-hoc onetailed t-test with WT, \*\**P* = 0.003). (d) Male DKO weanlings are visibly smaller and less developed than SKO and WT. Growth curves are provided in Supplemental 2).

With respect to the offspring produced during the breeding trials in this report, we found that the sex ratio of homozygous DKO offspring was skewed (0.85 male:female) but did not deviate significantly from the expected equal ratio (***χ***2 test, *n*=243, *P* = 0.22), and that the average DKO litter size was the same as WT mice (two-tailed t-test, n=4, *P* = 0.5). We also observed that unsexed 2-day old DKO pups from these same litters weighed significantly less than WT pups (Figure 3b, n=4 litters per strain, *P* = 0.004) and that DKO male weanlings weighed significantly less than WT (Figure 3c-d, n=21,45, *P* = 0.003; Figure S2, Additional File 2). Taken together, results from these studies suggest that loss of both PAD2 and PAD4 suppresses fertility and affects offspring weight. Previous studies have shown that pup weight is also commonly reduced in sub-fertile mouse strains, including androgen receptor knockout (ARKO) mice (36) which is relevant to the research reported here. Given the growing links between PADs and hormone signaling, we predicted that the effect of DKO on both fertility and pup weight was due to disrupted hormone signaling in these mice.

### Effect of Padi2/Padi4 DKO on pubertal onset

Pubertal onset is thought to be primarily driven by increased testosterone production in males and one well-established external marker of pubertal development in rodents is preputial separation (PS) (25,37,38). Therefore, as a further test of the hypothesis that hormone signaling is disrupted in DKO males, we next documented the timing of PS. Results show that DKO males took an average of 3.6 days longer than WT or SKO males to undergo PS (31.2 vs 27.6 days: ANOVA, *P* < 0.0001, Figure 4a,b). Given that the timing of pubertal onset has been repeatedly, yet inconsistently, linked with obesity (39,40), and been shown to differ between sexes within the same species/strain (41), we asked whether the delayed puberty in DKO males could be a side-effect of their smaller size at weaning. This does not appear to be the case, as DKO males are significantly heavier than WT and SKO at time of PS (ANOVA, *P* = 0.0002, Figure 4c), and the mean weight of males, regardless of strain, was the same at day 27, which is the average day of PS in WT and SKO (ANOVA, *P* < 0.0001, Figure 4d). Preputial separation is known to be an androgen-dependent process (38,42), strongly linked to hormone signaling along the hypothalamic-pituitary-gonadal (HPG) axis (43,44). These results further support the hypothesis that hormone signaling may be altered in DKO males. Additionally, these observations suggest that PAD2 and PAD4 play an important role in androgen-driven sexual development in males.

**Figure 4.**
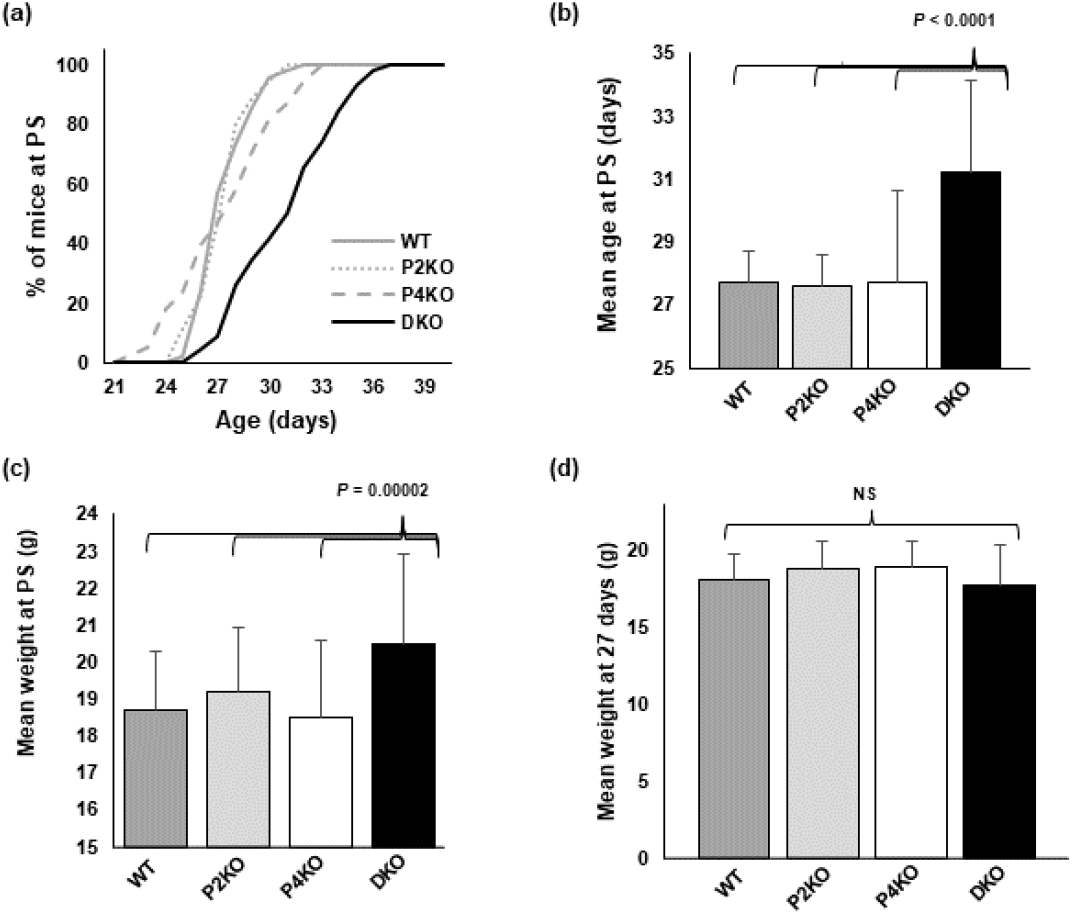
Pubertal onset is delayed in *Padi2/Padi4* DKO male mice. (a) Preputial separation occurred later in DKO males than WT and SKO strains, with (b) the mean age of PS significantly higher (n=49, 35, 38,46; ANOVA, *P* < 0.0001). (c) Smaller body size at weaning does not appear to be a factor, as DKO mice are significantly larger when they reach PS than all other strains (ANOVA, *P* = 0.0002), and (d) there is no significant difference in body weight between strains at day 27, the mean day of PS for WT and SKO mice (ANOVA, *P* = 0.5).

### Effect of Padi2/Padi4 DKO on serum hormone levels

As a more direct test of the hypothesis that hormone levels are altered in DKO mice, we next measured serum testosterone (T), luteinizing hormone (LH), and follicle stimulating hormone (FSH) in WT and DKO males. Results showed that levels of T, LH, and FSH differed significantly between DKO and WT at multiple timepoints (n=5 per time point). Although highly variable, serum testosterone levels were lower in DKO males than WT males at all time-points, most significantly at day 48 (*P* = 0.05, Figure 5). In contrast, FSH and LH levels were statistically similar between DKO and WT at the earlier time points, but at day 50 both hormones were significantly higher in DKO sera (*P* < 0.03). Although it is difficult to infer signaling responses from this data due to use of serum from different individual mice at each timepoint rather than sequential levels from the same mice across time,we were encouraged by the observation that serum T levels were consistently lower in DKO mice regardless of age, and that LH and FSH differed significantly between strains. Importantly, studies with ARKO mice also found that serum T levels are lower in mutant mice compared to WT mice (36). Given this observation, our findings lend support to the prediction that PAD enzymes play a role in androgen signaling *in vivo*.

**Figure 5.**
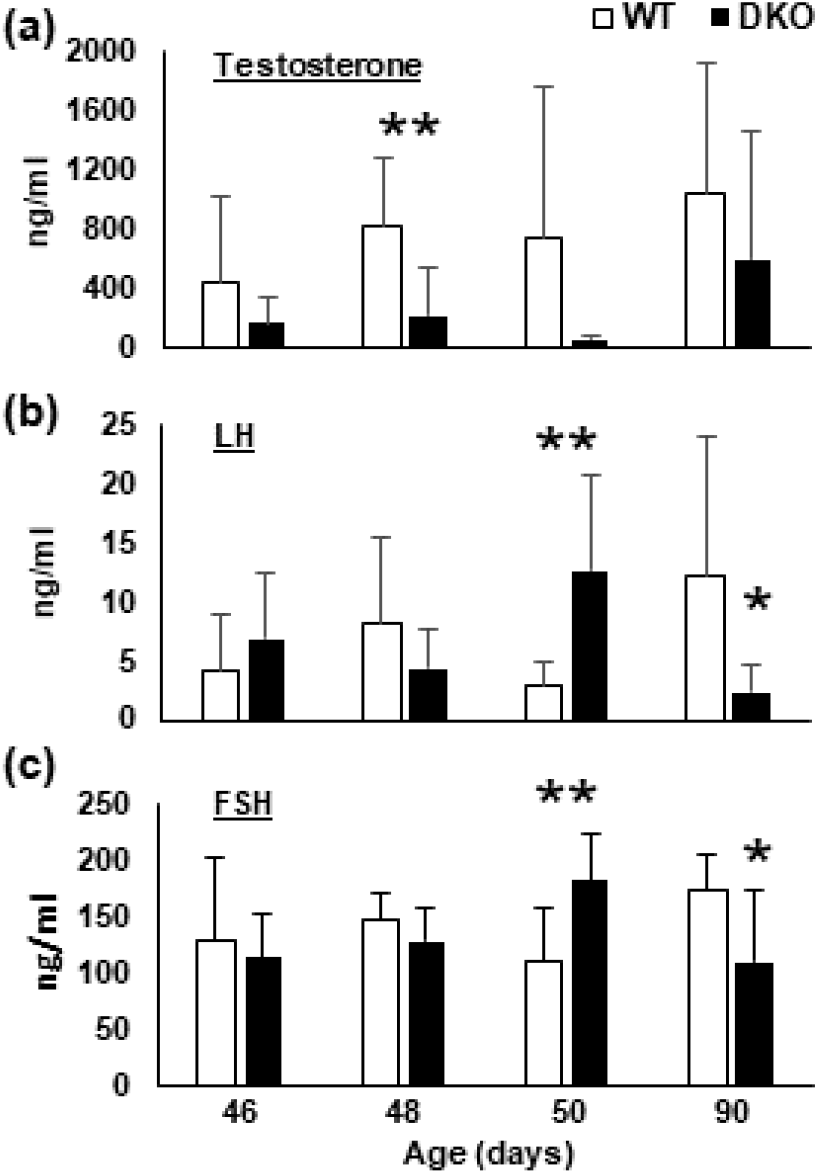
*Padi2/Padi4* DKO male mice show altered serum hormone levels. (a) Mean serum Testosterone levels were lower in DKO males when compared to WT males at all timepoints. (b.c) LH and FSH levels were significantly higher in 50-day old DKO mice, but significantly lower than WT in 90day old mice(n=5. one-tailed t-test, \**P*= 0.05, \*\**P* = 0.02.).

### Effect of Padi2/Padi4 DKO on testis size and histology

#### Testis Size

Hormone signaling within the HPG axis not only regulates pubertal onset, but also regulates body growth (41) and testis size (45,46). As a further test for potential associations between PADs and hormone signaling, we next investigated whether the total body weight and testis weight was altered in 90-day old adult DKO mice. Results showed that, while there was no difference in total body weight between the WT (m=27.4g) and DKO (m=27.4g, n=5,2-tailed t-test, *P* = 1), both absolute testis weight (*P* = 0.002) and the gonadosomatic index (testis weight as a percent of body weight) (*P* = 0.037) were significantly lower in DKO mice (Figure 6a,b). Yeh et al found that both pup size and testis weight are reduced in AR knockout mice (36), and our finding that DKO testes are significantly smaller than WT testes in adults fits well with the hypothesis that PADs play a role in androgen signaling in males.

**Figure 6.**
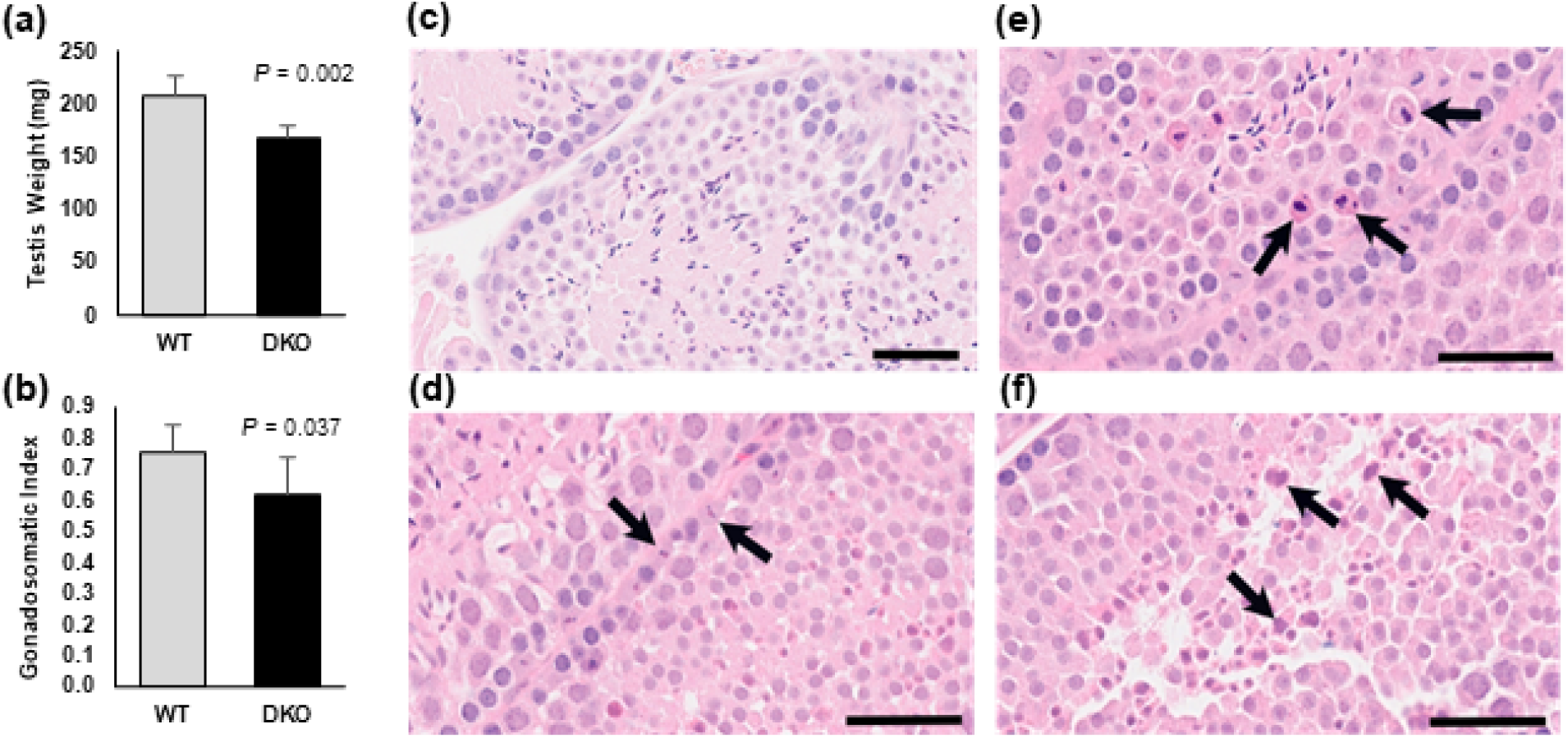
*Padi2/Padi4* DKO male mice have disrupted testicular development. In comparisons of 90-day old DKO to WT mice. DKO have (a) significantly smaller testes (n=5. one-tailed t-test, \**P*= 0.002) and (b) significantly lower gonadosomatic index (testes weight as% of body weight; n=5. one-tailed t-test, \*\**P*= 0.037). (c) Hematoxylin and Eosin stained sections of WT testis showing normal spermatogenic cell structures. (d-f) Hematoxylin and Eosin stained sections of DKO testis. Black arrows indicate: (d) apoptotic spermatogonia. (e) apoptotic spermatocytesand (f) atypical residual bodies (scalebars = 50*μ*m).

#### Testis Histology

The observation that DKO testes are smaller than WT testes raised the possibility that testicular histology may also be defective in DKO males. To test this hypothesis, FFPE sections of adult testes were stained with H&E and scored for cellular morphology by a board-certified pathologist (Table S1, Additional File 3). Results of this analysis found that there were variable degrees of germ cell apoptosis in the DKO testes when compared to WT testes, with spermatogonia and spermatocytes being the most significantly affected (Figure 6c-f). More specifically, apoptosis (mild/moderate grade) in spermatocytes (Figure 6d) was noted in 5 out of 5 DKO mice compared to only 2 out of 5 WT samples. Apoptosis (mild grade) of spermatogonia (Figure 6e) was noted in 4 out of 5 DKO mice testes compared to only 1 out of 4 WT mice. All five DKO testes also displayed moderate grade residual bodies (Figure 6f) when compared with WT testes, which only displayed mild grade residual bodies. Sertoli, Leydig cells and interstitium were overall unremarkable. The increased rate of apoptosis and residual bodies did not appear to result in a decrease in sperm production with the lumen of most tubules being lined with histomorphologically unremarkable elongated spermatids and spermatozoa. Interestingly, in ARKO mice, spermatogenesis was arrested at the pachytene stage of meiosis and there was an increase in apoptotic-like bodies within the tubules (36) suggesting that androgen signaling is critical for normal spermatogenesis. Taken together, these results support the hypothesis that PAD2/4 signaling may play a role in androgen signaling-mediated spermatogenesis.

### Effect of Padi2/Padi4 DKO on gene expression in the testis

In order to begin investigating whether PADs are associated with specific signaling pathways in the testis, and to more directly test the hypothesis that PAD2/4 play a role in regulating AR target gene expression, we next carried out RNA-Seq analysis of adult WT, P2KO, and DKO testes. Results show that 22 genes were upregulated and 19 genes were downregulated in P2KO samples when compared to WT samples (FDR cut-off 0.05, *p-adj* < 0.05). Additionally, we found that 263 genes were upregulated and 140 genes were down-regulated in the DKO samples compared to WT samples (FDR cut-off 0.05, *p-adj* < 0.05). Seventeen of the 22 genes that were significantly upregulated in the P2KO samples were also upregulated in the DKO samples. A comparison of upregulated genes is shown in Figure 7a, with the complete lists of differentially expressed genes (DEGs) provided in Tables S2 andS3 (Additional Files 4 and 5). Two hundred and forty-six genes that were uniquely upregulated in DKO testes were then subjected to DAVID gene ontology analysis. Of the 246 genes, 238 were identified in the mouse database and these genes were categorized into three domains: Cellular Components, Molecular Functions, and Biological Processes (Table S4, Additional File 6). Analysis of the Biological Processes domain finds that a majority of the DEGs are involved in reproductive functions (Figure 7b), thus supporting the phenotypic traits we observed. In addition, analysis of the Molecular Function domain finds that nearly all of the DEGs appear to be involved in either nucleic acid binding or protein phosphorylation (Figure 7c), thus further supporting the hypothesis that PADs play a role in regulating gene transcription. We next generated a hierarchical dendrogram and heatmap showing the top 100 most significantly up- and downregulated genes using summary counts over all pairwise tests of WT, P2KO, and DKO samples (FDR cut-off 0.05, *p-adj* < 0.05; Figure 8; Table S5, Additional File 7). Results show that there is a good correlation in gene expression levels between replicates for each experimental group. Additionally, the gene expression profile for the DKO samples is distinctly different from that of the WT samples, while there are some overlapping expression patterns when comparing the DKO and P2KO samples. We next narrowed this pool down to 41 transcripts that showed at least a 2-fold difference between strains for further investigation of their known activities. For this analysis, we used The Jackson Laboratory’s Mouse Genome Informatics (MGI) databases(47,48) and we reveiwed the current literature. Of these 41 transcripts, 15 were identified as lncRNAs, 20 were from protein coding genes, and 6 were undescribed or pseudogenes.

**Figure 7.**
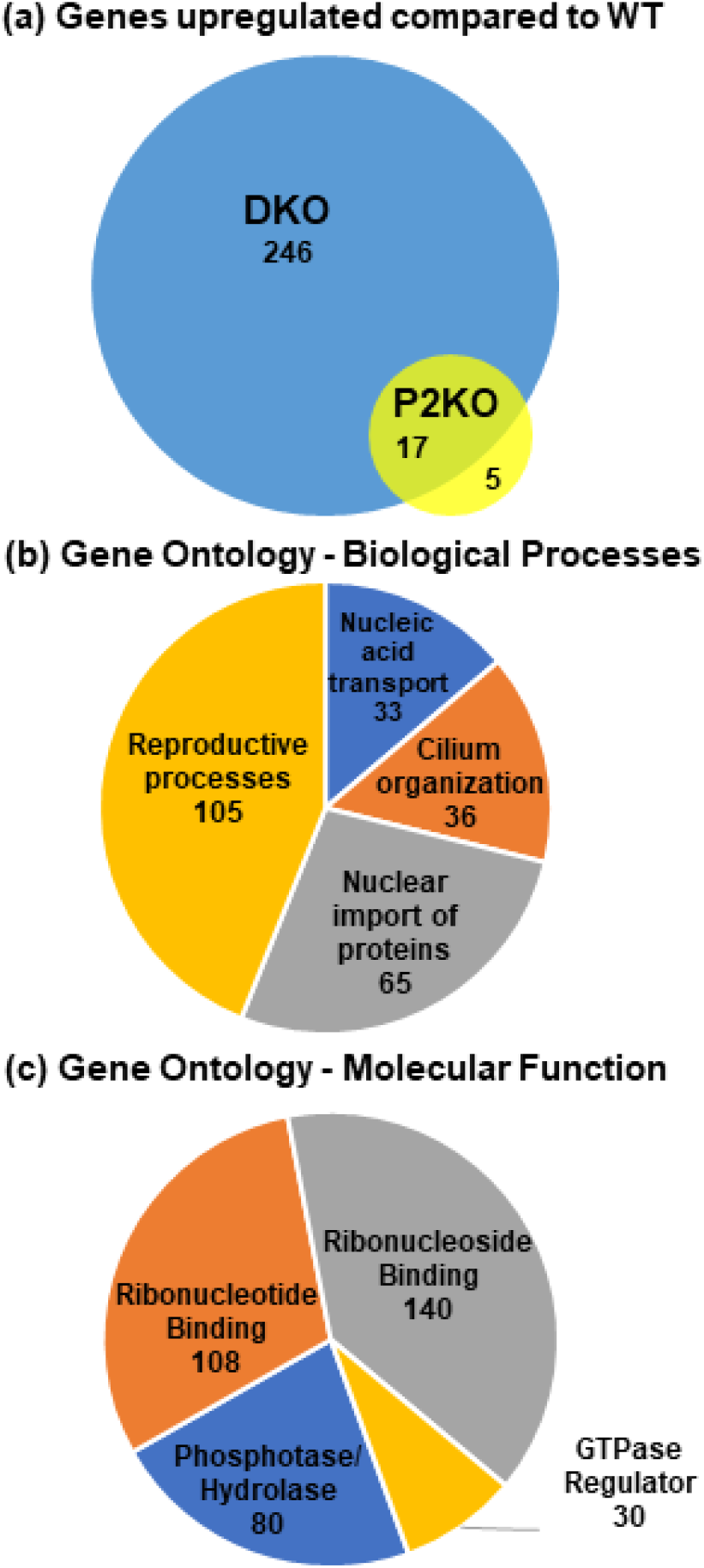
RNA-seq analysis of gene expression in 4-month old WT and DKO testes. (a) Venn diagram of differential gene expression shows that 263 genes are significantly upregulated in DKO in comparison to WT, with only 17 genes overlapping the P2KO profile. Significance was set as FDR < 0.05 and *p-adj* < 0.05. Gene lists are provided in Supplementals 4 and 5. (b-c) Gene ontology analysis with DAVID 6.8 divides these upregulated genes into (b) Biological Processes with reproductive pathways heavily represented and (c) Molecular Functions almost entirely represented by genes involved in nucleic acid regulation.

**Figure 8.**
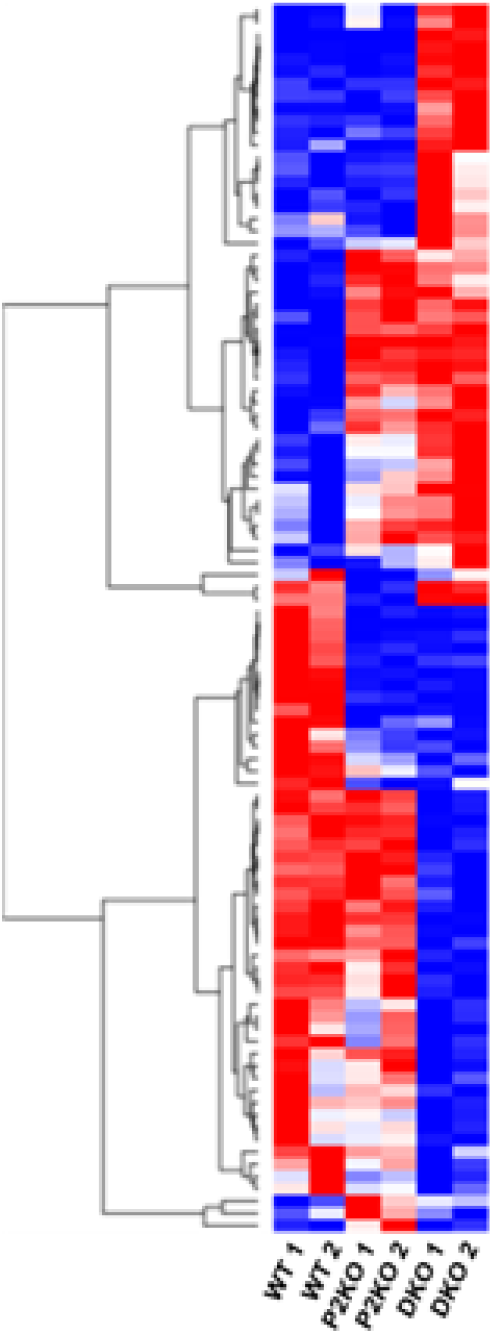
Heatmap and unsupervised hierarchical dendrogram illustrating the top 100 differentially expressed genes between 4-month old WT, P2KO and DKO testes. Significance was set as FDR < 0.05 and p-adj < 0.05. Red=upregulated, BI ue=down regulated. The list of DEGs used to create the heatmap is provided in Supplemental 7.

lncRNAs are generally known to be involved in transcriptional regulation through either binding DNA directly or changing methylation patterns on chromatin. Alternatively lncRNAs can directly interact with other chromatin modifiers to recruit or deflect their action on transcription (49,50). One lncRNA worth mentioning here, *Snhg12*, has been linked to regulation of the cell cycle and promotion of the Epithelial-Mesenchymal Transition in embryogenesis (51). In addition, Snhg12 has been shown to promote cell proliferation and inhibit apoptosis in multiple cancers (52), including testosterone-sensitive prostate cancer (53). Although we do not propose a mechanism, the almost 3-fold decrease in *Snhg12* levels in our DKO mice suggests loss of this factor may have played a significant role in the increased apoptosis and atypical residual bodies that are seen in Figure 6.

The 20 protein coding genes showing more than a 2-fold change in expression between DKO and WT included multiple members of two important gene families. First, the Kallikreins are a family of serine peptidases, many of which have been linked to AR expression and progression of androgen sensitive prostate cancer cells (54). Perhaps the best-known member of this family is prostate-specific antigen (KLK3), whose expression is mainly induced by androgen and is transcriptionally regulated by the androgen receptor (55). Additionally, the kallikrein gene locus is highly responsive to steroid hormones, having at least 14 functional HREs in this region. In fact, many researchers use kallikreins as markers of hormone receptor activity (56). In our DKO mice, *Klk21, Klk24* and *Klk27* are increased 3-fold compared to WT, and these three kallikreins have been shown to be expressed exclusively in Leydig cells and be responsive to testosterone(57). Additionally, these 3 kallikreins are upregulated in the FOXa3 KO mouse model which shows severe testicular degeneration, increased gonadal apoptosis and decreased fertility (57). The second over-represented group in our DKO mice are Zinc-finger proteins, a diverse family generally known as regulators of gene expression. Six Zinc-finger proteins are differentially expressed in our DKO mouse testes, two of which are of particular interest here. *Zfp982* has previously been shown to be involved in cell lineage differentiation in embryogenesis (58) and the 12-fold decrease in levels of this factor in our DKO mice may have significantly disrupted developmental processes. *Zfp979* (also known as *Ssm1b*) initiates DNA methylation and inhibits transcription, in undifferentiated embryonic stem cells as well as in adult murine germ cells (59). The 20-fold decrease of *Ssm1b* seen in our DKO testes could allow expression of a multitude of proteins detrimental to early development and germ cell maturation.

Within the remaining differentially expressed coding genes, several stand out as relevant to our discussion for their potential roles in steroid signaling and the HPG axis. *Runx3* is decreased 6-fold in our DKO testes. This this factor has been shown to regulate steroidogenesis and gonadal development in female mice (60). Whether it acts similarly in male mice is currently unknown. Mutations in *Spry4*, which is upregulated 3-fold in our DKO testes, has been implicated in incomplete sexual maturation and infertility due to decreased GnRH activity in human males (61). *Sdc3* (upregulated 12-fold) is considered a regulator of obesity and has been proposed as a modulator of gonadal steroid function (62). And finally, altered levels of *Mrto4* in Balb/C mice (upregulated 3 fold in our DKO) have been linked to disruptions in spermatogenesis, fertility and testosterone levels in mice (63). These findings further support our hypothesis that PADs play an important role in regulating gene expression in the testis, and more specifically, in regulating AR-mediated gene expression.

## CONCLUSIONS

Our previous work with PAD2 and PAD4 in breast cancer cell lines has found that both of these PAD isozymes appear to regulate ER target gene expression via citrullination of histone residues at ER binding sites. More recent studies have shown that PAD2 also appears to facilitate androgen receptor target gene expression in prostate cancer cells via similar mechanisms. Together, these observations suggest that PAD2 and PAD4 may represent key mediators of hormone signaling in mammals. While these *in vitro* findings are intriguing, an important next step in linking PADs with hormone signaling is demonstrating that the phenomenon also occurs at the organismal level. While our *Padi2* and *Padi4* SKO mouse models have failed to show a reproductive phenotype, outcomes from our *Padi2/Padi4* DKO studies provide a series of clear associations between PADs and hormone signaling and, to our knowledge, this study is the first to make such genetic links using animal models.

We predict that the phenotypic effects of DKO that we have observed in this study are due to the lack of PAD2 and PAD4 activity in the testis. Our DKO mice described here exhibit several phenotypic traits (delayed puberty, small offspring and decreased serum testosterone) that are characteristic of other sub-fertile strains, including a number of AR knockout mouse models (64). Since PAD2 and PAD4 have previously been implicated in interactions with estrogen and androgen receptors, it is not unexpected to see reproductive defects in these mice. However, our additional finding of abnormal serum testosterone, LH and FSH levels indicates a wider reaching disruption of the HPG axis with implications for more than just reproductive fitness.

## LIST OF ABREVIATIONS

Padi or PAD: Peptidylarginine deiminase
DKO: Double knockout
SKO: Single knockout
WT: Wildtype (FVB/NJ)
E2: Estrogen
ER: Estrogen receptor
ERE: Estrogen response element
AR: Androgen receptor
ARKO: Androgen recepter knockout
PS: Preputial separation
HPG: Hypothalamic-pituitary-gonadal axis
T: Testosterone
FSH: Follicle stimulating hormone
LH: Lutenizing hormone
H&E: Hematoxylin and Eosin stain
DEGs: Differentially expressed genes

## DECLARATIONS

### Ethics approval and consent to participate

All animal procedures were approved by Cornell University’s Animal Care and Use Committee (IACUC protocol #2007-0115) and are in compliance with the US National Research Council’s <u>Guide for the Care and Use of Laboratory Animals</u>, the US Public Health Service’s <u>Policy on Humane Care and Use of Laboratory Animals</u>, and <u>Guide for the Care and Use of Laboratory Animals</u>.

### Consent for publication

Not applicable.

### Availability of data and materials

The datasets supporting the conclusions of this article will be available upon publication in NCBI’s Gene Expression Omnibus (GEO) repository (GSE194034, https://www.ncbi.nlm.nih.gov/geo/). *Padi2/Padi4* double knockout mice are available from the corresponding author upon reasonable request, and in accordance with Cornell University’s Material Transfer Agreement.

### Competing interests

All authors declare that they have no competing interests.

### Funding

This research did not receive any specific grant from any funding agency in the public, commercial or not-for-profit sector.

### Authors’ contributions

Experimentation was conceived and conducted by KLS, C Mukai, BAM and SAC. Breeding and phenotype analysis by KLS and SAC. Histological scoring by EAD and SN. RNA sequencing and analysis by JKG, AET, and FA, with visualization by C Mittal and KLS. All authors read and approved the final manuscript.

## Acknowledgements

The authors acknowledge the excellent skills and technology provided by Cornell’s Stem Cell & Transgenic Core Facility to create our double knockout mice. We also thank the Baker Institute husbandry technicians for the outstanding care of the animals used in this research.

## ADDITIONAL FILES

**Additional File_1.pdf**

Figure S1. Full PCR gels and membranes for authentication of DKO mice.

**Additional File_2.pdf**

Figure S2: Growth curves of WT, SKO and DKO mice

**Additional File_3.pdf**

Table S1. Histological scoring of cellular abnormalities in 90-day old WT and DKO testes.

**Additional File_4.xlsx**

Table S2. Significantly differentially expressed genes in 4-month old DKO testes compared to WT (FDR < 0.05 and Benjamini-Hochberg *p-adj* < 0.05).

**Additional File_5.xlsx**

Table S3. Significantly differentially expressed genes in 4-month old P2KO testes compared to WT (FDR < 0.05 and Benjamini-Hochberg *p-adj* < 0.05).

**Additional File_6.xlsx**

Table S4. DAVID results of gene ontology categories over-represented in 4-month old DKO testes compared to WT.

**Additional File_7.xlsx**

Table S5. Top 100 most up- and downregulated genes from 4-month old testes (FDR < 0.05 and Benjamini-Hochberg *p-adj* < 0.05), ranked by Fold Change and used to create the heatmap in Figure 8.

## Notes

### Competing Interest Statement

The authors have declared no competing interest.

### Summary of Updates

Added data for single knockout mouse strains and clarified breeding trial strategies. Showed validation that Pad1 and Pad3 are not disrupted. Changed title and text to be more descriptive and decrease assumptions. Added more discussion of potentially important genes that are differentially expressed in the DKO testes.

