## Supplementary figures and images for "Delayed puberty, gonadotropin abnormalities and subfertility in male *Padi2/Padi4* double knockout mice"

### Additional File 1

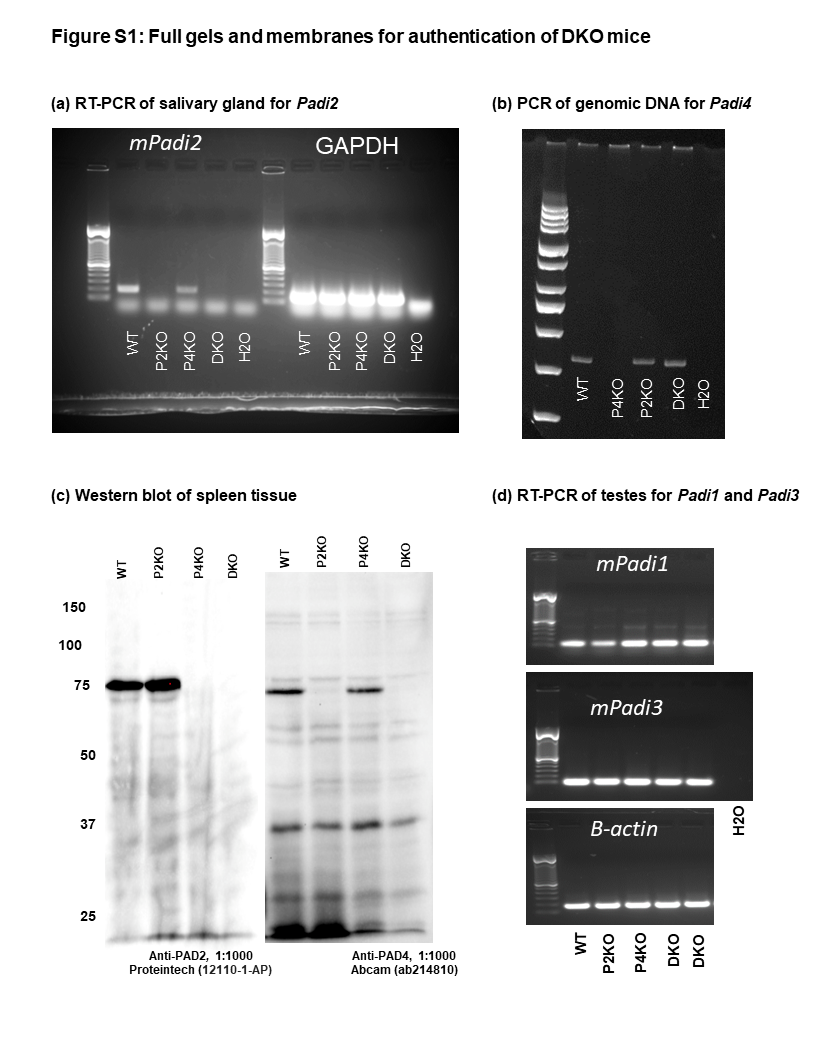

### Additional File 2

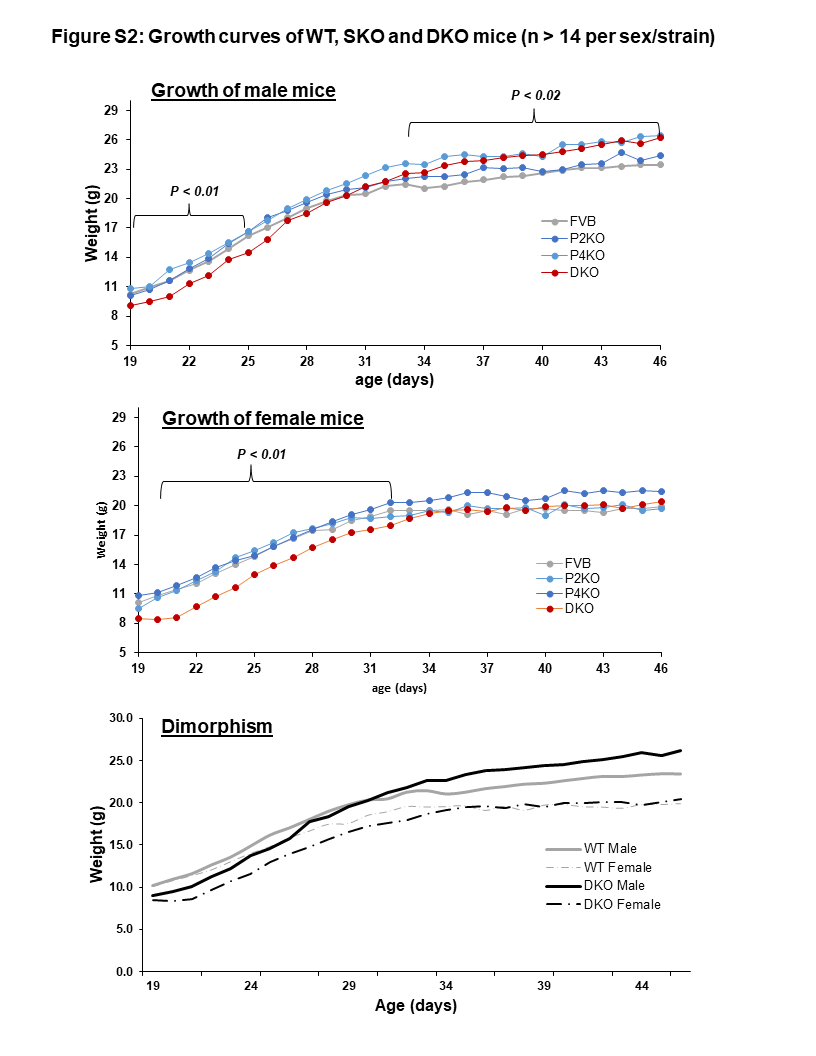
